# The art of snacking: Innovative food extraction in a synanthropic species is associated with food exposure and food categorization

**DOI:** 10.1101/2021.09.01.458608

**Authors:** Tejeshwar Dhananjaya, Sayantan Das, Amal K. Vyas, Prakhar Gahlot, Mewa Singh

**Author notes:** School of Biology, Indian Institute of Science Education and Research, Thiruvananthapuram-695551, India (Amal K. Vyas). Department of Biological Sciences, University of New Orleans, Los Angeles-70148, USA. **Correspondence:** Sayantan Das, Biopsychology laboratory, Department of Psychology, Institute of Excellence, University of Mysore, Manasagagothri, Mysuru-570006, India.

## Abstract

Extractive foraging is generally studied from the perspective of behavioral flexibility, cognitive ability, innovation and social learning. Despite its potential to elucidate synanthropic adaptation in species exploiting enclosed anthropogenic food, research on extractive foraging under urban conditions is limited. Since a large extent of anthropogenic food is packaged and contains highly processed food, processes of identification/extraction of food by nonhuman species become intriguing themes of research. We studied how processing status of embedded food determined extraction decisions across groups of a species differing in exposure and familiarity to the food. Further, we tested the generalizability of extraction methods. Experimenting with wild bonnet macaques (Macaca radiata), we found exposure- and form (native/shelled/peeled)-specific familiarity to peanuts, state (raw/boiled/roasted)-specific distinction in depeeling, and exposure- and state-specific differences in methods of depeeling. Group with the highest exposure to peanut differed in its propensity to use sophisticated extraction methods, e.g. depeeling by rubbing between palms (bimanual asymmetric action) and rubbing against horizontal substrata (unimanual action). The innovative methods were also extended to roasted peas and chickpeas by the urban group. Our study establishes a causal relationship between familiarity and processing status of food and shows the generalized extension of extraction methods based on food categorization.

**Summary Statement:** Nonhuman species in cities face upheaval challenges of accessing enclosed and highly processed anthropogenic food. We studied the effects of minor processing of enclosed food on its extraction decisions.

## INTRODUCTION

Modern human diet features an extraordinary variety of processed foods containing primary ingredients (e.g. grain, pulse and nut) that are synthesized through a series of physiochemical processes (see Fellows, 2009) such that the original ingredients are nearly impossible to cognize in the final product. Though primary food processing is necessary to ensure edibleness and palatability of many food items (Dwyer et al., 2012; Keding et al., 2013), artificial processes alter the nascent properties of the food in ways that are unparalleled under natural conditions. Processing, including amalgamation of a large variety of primary food items alters appearance, texture, taste, smell and mouth feel (Davidou et al., 2020; Fardet & Rock, 2019) of ingredients rendering their acceptance as food unapparent. Deviation from naturalness (Rozin, 2005; Scott & Rozin, 2017) can trigger the perception of food as hazardous (Scott et al., 2018) and less acceptable (Gaskell et al., 2011), often by evoking dread (Slovic, 1987) and disgust (Egolf et al., 2019; Siegrist et al., 2018; Verbeke et al., 2015). Choice of processed food is thus, stimulated by the ideation of naturalness, which is mediated by an “instinctive desire (biophilia) for the experience of ancestral environment” (Rozin, 2005).

Processed foods often spill over directly or indirectly as allochthonous materials and assimilate in the environment of synanthropic nonhuman species through voluntary/involuntary (e.g. spilling) human provisioning and by improper waste disposal (Newsome et al., 2015; Oro et al., 2013), respectively leading to a sustained influx of novel anthropogenic food sources (Becker & Hall, 2014; Tuomainen & Candolin, 2011). Consequently, once exposed, few synanthropic species could proactively explore sources of processed food, like anthropogenic spaces (e.g. kitchen, restaurant/bakery, campsite, open fruit/vegetable market, vehicles) and artificial containers (e.g. garbage/rubbish bins, picnic/food basket, bird/pet feeders, carry bag) that alternately enhance chances of encountering novel anthropogenic food items (Calle & Gawlik, 2011; Osterback et al., 2015). Plaza and Lambertucci (2017) found evidence of 98 vertebrate species (amphibian-5, reptile-5, bird-54, mammal-34) being present in garbage dumps. Among synanthropic mammals, generalist nonhuman primates (primates subsequently) are epitomes of anthropogenic resource exploiters (Fuentes, 2012; Priston & McLennan, 2013) despite a robust neophobic system (e.g. Johnson, 2000). While crop foraging allows primates to integrate novel cultivars into their diets (see Hill, 2018), processed food is more widely available beyond agricultural landscapes. Processed food is encountered at open/closed dining sites (e.g. soft drinks-vervet monkey: Else, 1991; instant snacks and potato crisp-long-tailed macaque: Sha et al., 2009), waste disposal spaces (e.g. cooked food, jam, baked items and milk products-yellow baboon: Altmann & Muruthi, 1988; fast food, cooked rice, noodles, cooked fish and cooked meat-long-tailed macaque: Sha & Hanya, 2013), food storage spaces (e.g. bread, sugar and candy-chacma baboon: Kaplan et al., 2011; butter popcorn-lion-tailed macaque: Shanmughanandam, 2019), animal enclosures (ostrich pellet-chacma baboon: Doorn et al., 2010; maize porridge-vervet monkey: Saj et al., 1999), during human provisioning (e.g. Indian fried bread, Indian roasted bread, icecream, roasted gram and roasted peanut-rhesus macaques: Wolfe, 1992; icecream, chips, cooked spicy rice and wheat preparations-bonnet macaque, Marty et al., 2020) and during proximate interactions with unsuspecting people (e.g. icecream cone-rhesus macaque: Wolfe, 2002; biscuits, instant snacks and soft drinks-bonnet macaque: Mangalam & Singh, 2013). Many commercial food items require extraction from their synthetic packaging, which often obfuscates sensory cues necessary for their identification, edibility and/or nutritional quality. Under such circumstances, how certain species learn to identify food items from their packaging (sealed foil, jar, can, tetrapack, bottle, poly bag, etc.) and succeed or fail in extracting them are intriguing themes of research within urban foraging in a proximate sense, and urban adaptation in an ultimate sense (see Mangalam & Singh, 2013). Identification and extraction of processed food resources are thus, ubiquitous problems for urban primates. It is essential to acknowledge though that empirical investigation of processed food extraction by synanthropic primates has critical ethical concerns generally and specifically, when they subsist under perilous and hostile anthropogenic conditions.

Most studies on extractive foraging in primates have either documented extractive foraging in naturalistic settings with natural food items (e.g. shellfish cracking in long-tailed monkey, Gumert & Malaivijitnond, 2012; walnut ‘processing’ in Japanese macaques, Tamura, 2020; plant underground storage organ ‘processing’ in capuchin, Truppa et al., 2019) or have employed artificial extraction tasks in captive (e.g. Crast et al., 2010; Kendal et al., 2015) and non-captive settings (e.g. Amici et al., 2020; Canteloup et al., 2020), most of which involved placing natural food items within strategically designed boxes. Although our understanding of behavioral diversity, manual dexterity, cognition, innovation, intelligence, individual/social learning and culture/traditions has emerged from the aforementioned studies, how processing status of enclosed food determines extraction decisions has not received ethological attention. An additional variable that appears to play a decisive role in the acquisition and development of extractive foraging behavior is exposure (to demonstrators and to elements associated with the foraging task), sometimes at a critical developmental phase (Inoue-Nakamura & Matsuzawa, 1997; Neadle et al., 2020). However, despite its broad role in individual/social learning (see Das et al., 2020; Hoppitt & Laland, 2013; Kummer & Goodall, 1985) and alternately, in catalyzing individual reinnovation of latent solutions (Reindl et al., 2018), the role of prior/extended exposure in execution of food extraction tasks has been largely biased towards technical (i.e. tool-use assisted) extractive foraging (e.g. Vale, Davis, Lambeth, et al., 2017; Van Schaik et al., 1999). As a result, we considered the roles of age, exposure (defined as rate of encounter), familiarity (defined as the ability to recognize/identify) and processed state (of food) on characteristic features of extractive foraging in our study.

We used 2 states of chemical processing, boiled and roasted and 2 forms of physical processing, shelled and peeled along with the unprocessed state (raw) and unprocessed form (native) to compare occurrence/method of extraction for a combined set of three forms by three states, resulting in nine combinations (cf. Canteloup et al., 2021). Since peeled peanuts do not require extraction, we used the peeled condition to compare familiarity. Unlike the aforementioned studies, this experimental design aided detailed investigation into extraction behavior, i.e. occurrence of shell/peel removal and methods of shell/peel removal. We chose bonnet macaques (Macaca radiata) as our study model owing to their high-density synanthropic subsistence from forest-agriculture matrix to temple/tourist sites (Singh, 2019), dietary diversity and flexible foraging skills (Deshpande et al., 2018; Mangalam & Singh, 2013). Three groups of bonnet macaques differing in their exposure to peanuts were chosen for the study. Incidentally, the groups also aligned along the rural-urban gradient. We hypothesized that (1) the group with a high exposure to peanuts has the most complex/optimal extraction method and (2) processing state of peanut influences choice of extraction and method of extraction. Based on the proposition of sensitive period for learning extractive foraging skills, we anticipated adults (>4 years; of the ‘High’ exposure group) to employ complex extractive methods more frequently than the young.

A strong enabling factor for synurbization (i.e. adaptation to urban conditions, Luniak, 2004; cf. Francis & Chadwick, 2012) is the ability to categorize objects (i.e. ‘perceive, discriminate and classify cues’, Barrett et al., 2019). Categorization can not only attenuate neophobia but facilitate an informed response (through adjustment of previous response) to unfamiliar objects that resemble previously encountered items (Barrett et al., 2019). We tested categorization among individuals of the ‘High’ exposure category since they were relatively the most urbanized. We chose roasted states of pea (seed) and chickpea and assessed whether they elicited extractive foraging responses similar to peanut since they have similar seed coat (i.e. testa) structures. Finally, since the exposure categories were also social groups, we expected individuals within a group to be homogeneous and individuals between groups to be heterogeneous with regard to their choice of extraction methods, a direct corollary of the theory of latent solutions (see Tennie et al., 2009).

## RESULTS

Based on experiments conducted in the ‘Introductory’ phase on 66 individuals across three groups, we obtained a total of 447 records, 1041 records and 607 records of native, shelled and peeled forms of peanut with a minimum of 35 putative representations of each state within each form and each exposure/group (Table 1). Demographic and population coverage of the sample set was extensive enough to capture intra-group variation in peanut extraction behavior. However, it is essential to note that partial habituation of TB necessitated bulk offer of peanuts in small clusters during four sessions. Similarly, our attempts to minimize social learning (of peanut extraction) failed when young juveniles (<2 years of age) accepted peanuts in isolation but preferred moving closer to peers during feeding. Since our coverage of young juveniles was low, we classified age into just two classes, young (<4 years) and adult (>4 years). The latency to accept peanuts emerged to be a highly exposure-specific response with all forms and states accepted by the three groups albeit with distinct latencies. When presented in native form, removal of pod largely proceeded in an identical manner (i.e. by biting with mouth while supporting peanut with one/both hand(s)) across exposure and hence, was excluded from analyses of group comparisons. Intriguingly, a single adult male in FH group deshelled peanuts by rubbing it against an artificial substratum (1 successful; 2 failed attempts). Extractions of the shelled form of peanut presented the maximum variation in occurrence and method of extraction and varied across state, and exposure *state and, and exposure and state, respectively. The most common peanut extractions were by oral and forelimbs-oral methods. Regardless, palm rubbing and rolling on substrata were recorded in the roasted-shelled peanuts almost exclusively in the ‘High’ exposure group. Finally, in the ‘categorization phase’ the innovative extraction methods were extended to roasted peas and roasted chickpeas.

**Table 1.** Sample size of the Introductory experiment based on form and state of peanuts with details on participants and outcomes. Extraction in context of native form is deshelling and in context of shelled form is depeeling.

| Peanut form | Peanut state | Number of peanuts recorded |  |  | Number of unique participants |  |  | Number of peanuts extracted |  |  | Number of peanuts ingested |  |  |
| --- | --- | --- | --- | --- | --- | --- | --- | --- | --- | --- | --- | --- | --- |
|  |  | BT | FH | TB | BT | FH | TB | BT | FH | TB | BT | FH | TB |
| Native | Raw | 41 | 80 | 35 | 11 | 29 | 12 | 41 | 80 | 35 | NA | NA | NA |
|  | Boiled | 40 | 46 | 41 | 11 | 25 | 13 | 40 | 46 | 41 | NA | NA | NA |
|  | Roasted | 58 | 55 | 50 | 15 | 28 | 16 | 58 | 55 | 50 | NA | NA | NA |
| *Total* |  | 139 | 181 | 127 |  |  |  | 139 | 181 | 127 |  |  |  |
| Shelled | Raw | 63 | 210 | 47 | 15 | 34 | 12 | 28 | 124 | 6 | 63 | 210 | 47 |
|  | Boiled | 57 | 142 | 77 | 14 | 24 | 12 | 57 | 101 | 69 | 57 | 142 | 77 |
|  | Roasted | 172 | 177 | 97 | 16 | 34 | 16 | 171 | 169 | 95 | 172 | 176* | 97 |
| *Total* |  | 292 | 529 | 221 |  |  |  | 256 | 394 | 170 |  |  |  |
| Peeled | Raw | 56 | 59 | 62 | 15 | 34 | 13 | NA | NA | NA | 56 | 59 | 62 |
|  | Boiled | 64 | 55 | 74 | 15 | 27 | 14 | NA | NA | NA | 64 | 55 | 74 |
|  | Roasted | 97 | 53 | 87 | 16 | 34 | 16 | NA | NA | NA | 97 | 53 | 87 |
| *Total* |  | 217 | 167 | 223 |  |  |  |  |  |  | 217 | 167 | 223 |

BT-Bull Temple group; FH-Foot Hill group; TB-Tirumakadul Betahalli group; \*In one case a peanut was accidentally dropped; NA-Not applicable

### Latency of peanut acceptance

Based on a two-way random effects model (ICC (2,2), see Koo & Li, 2016), the mean measure of inter-rater consistency in latency of accepting (LA) peanut was 0.83 (95% CI: 0.578, 0.917; F=7.96, p<0.0001). We used a subset of data to model latency and considered a total of 301 data points across 54 individuals for the analyses. The raw scores were transformed into integers by multiplying with a factor of 100 to perform data modeling. Controlling for the effects of repeated measure on individual macaques, we found LA of ‘Low’ category to be 3.4 times greater than ‘High’ category and 2.5 times greater than ‘Moderate’ category while LA of ‘High’ and ‘Low’ exposure categories were comparable (Table 2). In partial fulfillment of our hypothesis, LA differed across forms but not across states with interaction effects between exposures and forms, and exposures and states. While the ‘Low’ exposure category was the least familiar, the ‘Moderate’ exposure category was slightly unfamiliar and the ‘High’ exposure category was the most familiar with peeled form, the exposure categories were comparable in their familiarity to native and shelled forms (Table 2). With regard to peanut states, though the ‘Low’ exposure category was slightly unfamiliar with the boiled form, the other exposure categories did not distinguish among peanut states (see Fig. S1 for model diagnostics).

**Table 2.**
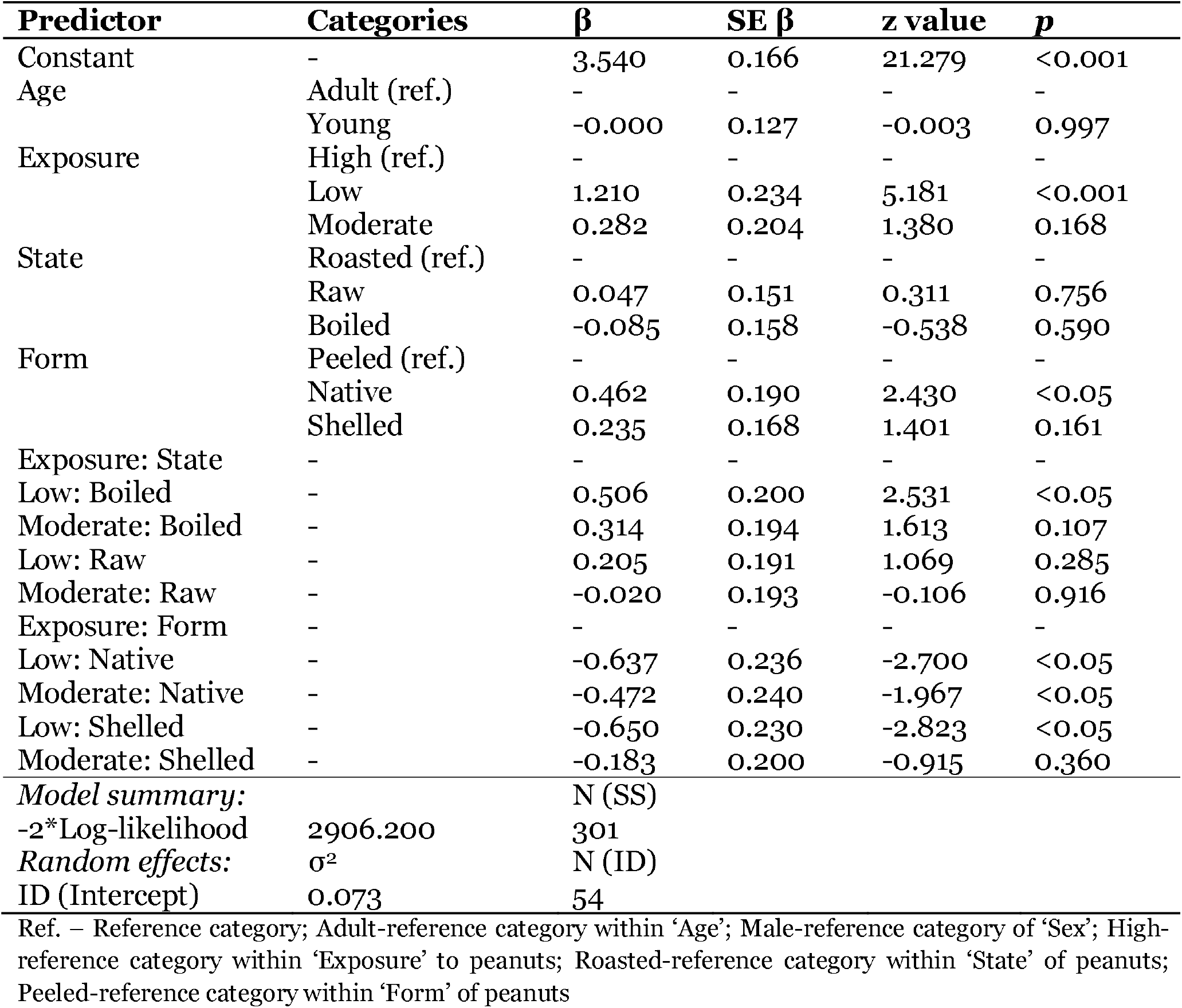
Results of the multilevel generalized linear model with ‘latency of acceptance’ as the dependent variable and age (Young and Adult) of participating macaque,exposure of group to peanuts (High, Moderate and Low), form of peanut (Native, Shelled and Peeled) and state of peanut (Raw, Boiled and Roasted) as predictor variables along with interaction effects of exposure and state of peanut. Latency was modeled as a negative binomial function (due to a positively skewed distribution) with log-transformed link function.

### Occurrence of testa removal within peanut extraction behavior

Inter-rater agreement in the occurrence of testa removal was 0.87 (Cohen’s κ, UL-LL: 0.84-0.91). The multilevel binomial logistic regression to model deskinning of peanuts showed that age was not a veritable predictor and hence, the factor was dropped from analysis (following Burnham & Anderson (2002)). In contrary to our hypotheses, deskinning decisions remained invariable across exposures and there was no interaction between exposure and state of peanuts (Table 3). However, raw state was less deskinned than roasted state by a factor of ∼700 in accordance to our expectation of state-specific extraction decisions (see Fig. S2 for model diagnostics).

**Table 3.**
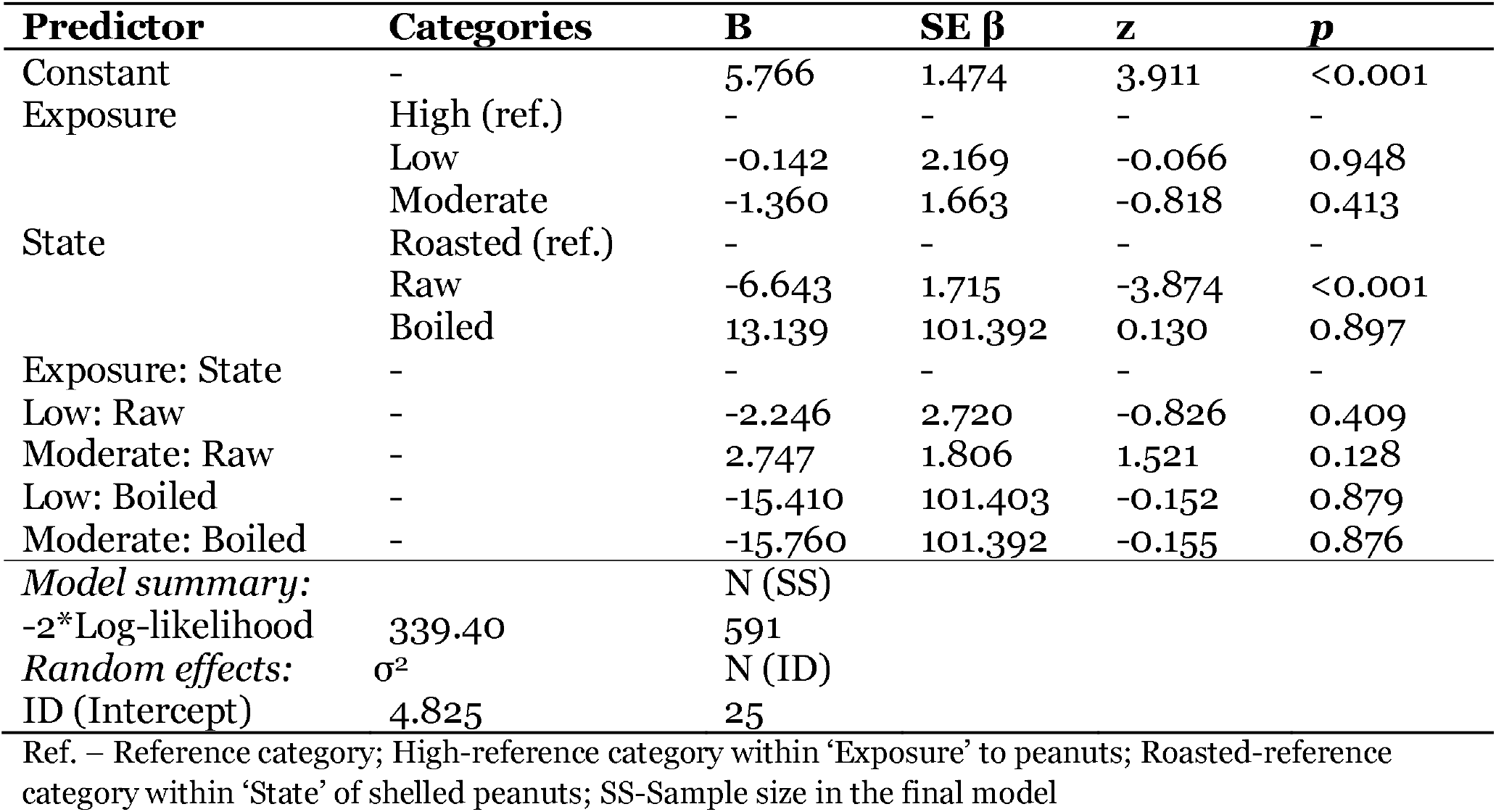
Model summary of the multilevel binary logistic regression to determine factors that affected the extraction of shelled peanut in the ‘Introductory’ experiment. The fixed factors hypothesized to affect extraction were exposure (High, Moderate and Low) of groups to peanuts and state of peanut (Raw, Boiled and Roasted) with probable interaction between exposure and state peanuts. Macaque identity (ID) was used as random factor and macaques with a representation of >15 cases were included in themodel. Occurrence of extraction was coded as ‘1’. The model was fit by maximum likelihood and used Laplace approxim

### Methods of testa removal within peanut extraction (Introductory experiment)

From the perspective of organ use, the primary modes of testa removal used either, the hands, the mouth or both, the hand and the mouth. Considering the organs and actions used to extract peanuts, we characterized extraction methods into (1) within mouth (M), wherein testa removal from shelled peanuts occurred within the mouth by using the tongue against the mouth palate (Supplementary Video MS1), (2) hand(s) and mouth (HM; unimanual and bimanual), which included nibbling with teeth while supporting with hand(s) (Supplementary Video MS2), (3) palm rub (PR; bimanual symmetric digit/hands action), wherein a peanut was rubbed between the palms (resulting in testa removal; Supplementary Video MS3) and finally, (4) rub on ground (RG; unimanual symmetric digits action), which involved rolling the peanut under the palm against a hard and horizontal substratum (Supplementary Video MS4). Inter-rater agreement in determination of extraction method was 0.92 (Cohen’s κ, UL-LL: 0.90-0.94). To enable controlling for the random effects of repeatedly sampling individual macaques in determining covariates of extraction methods, we divided the dataset into instances in which M and HM were used and instances in which PR and RG were used such that the methods could be modeled as 2 multilevel binary logistic regression models. From Figure 1, it is apparent that PR and RG were employed by ‘High’ exposure group and strictly in the shelled-roasted peanut condition. Only two instances of RG were recorded in ‘Moderate’ exposure category, once in shelled-raw peanut by a young adult male and the other in shelled-roasted peanut by an old adult male. Following Hegyi & Laczi (2015), the optimum regression model to determine covariates of HM included age, exposure and state of peanuts as determinants. Young macaques did not differ from adult macaques in use of HM for peanut extraction (Table 4). In affirmation to our hypothesis, the ‘High’ exposure group used HM 19 times more than the ‘Low’ exposure and nine times more than the ‘Moderate’ exposure groups, while the ‘Low’ exposure group remained comparable to ‘Moderate’ exposure group. Even in context to state of peanuts, HM was used to extract peanut in the raw state twice more than the roasted state though HM did not differ between roasted and boiled states and between raw and boiled states (Table 4; see Fig. S3 for model diagnostics).

**Table 4.**
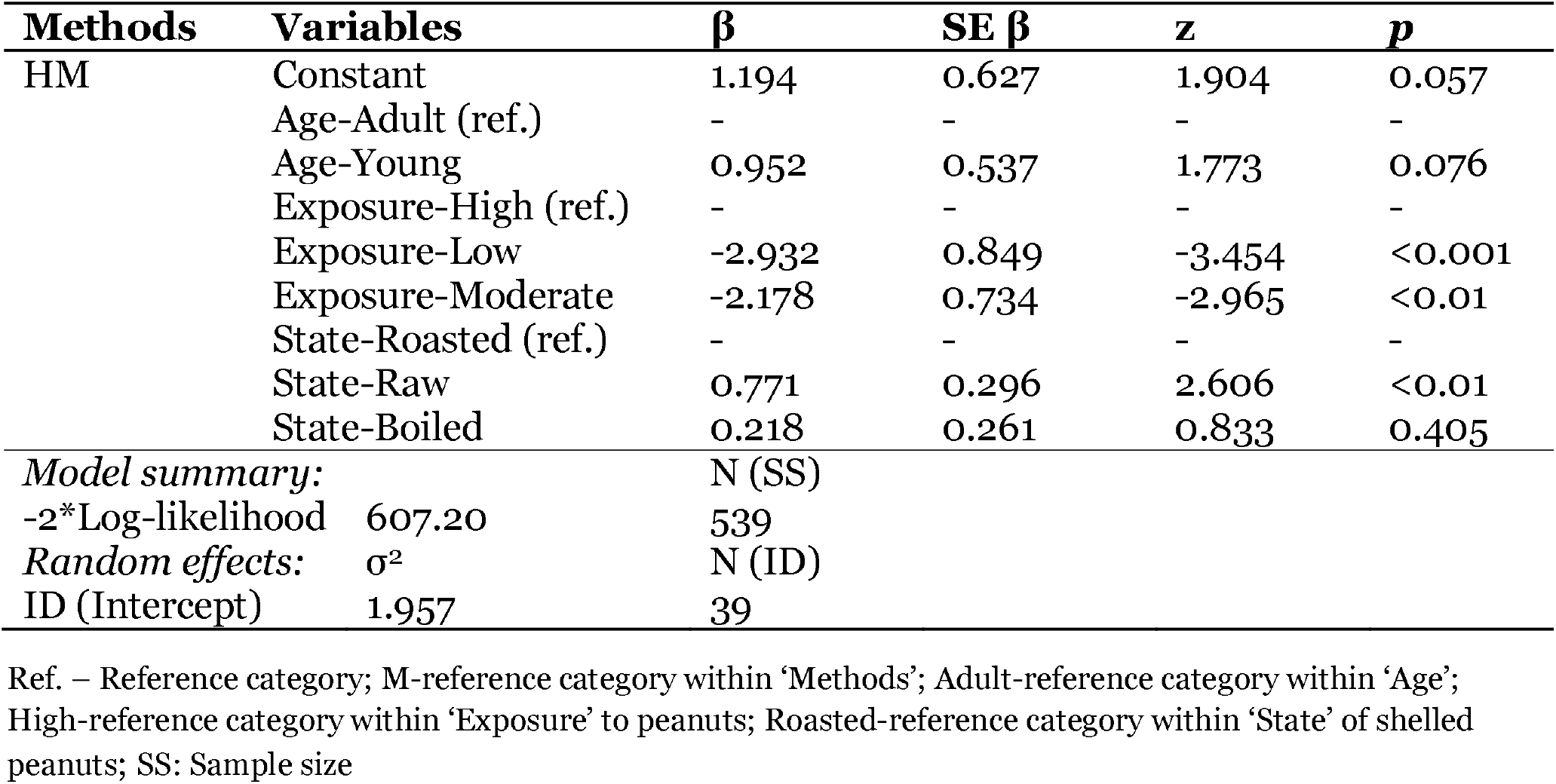
Model summary of the multilevel binomial logistic regression to identify determinants of the Hand-Mouth (HM) method of peanut extraction in the ‘Introductory’ experiment. The fixed factors were age (Young and Adult) of participant, exposure (High, Moderate and Low) to peanuts and state of shelled peanut (Raw, Boiled and Roasted) and an interaction effect between exposure and state of peanut was considered. The identity of individuals (ID) was used as random factor. Model fit used maximum likelihood with Laplace approximation.

**FIGURE 1.**
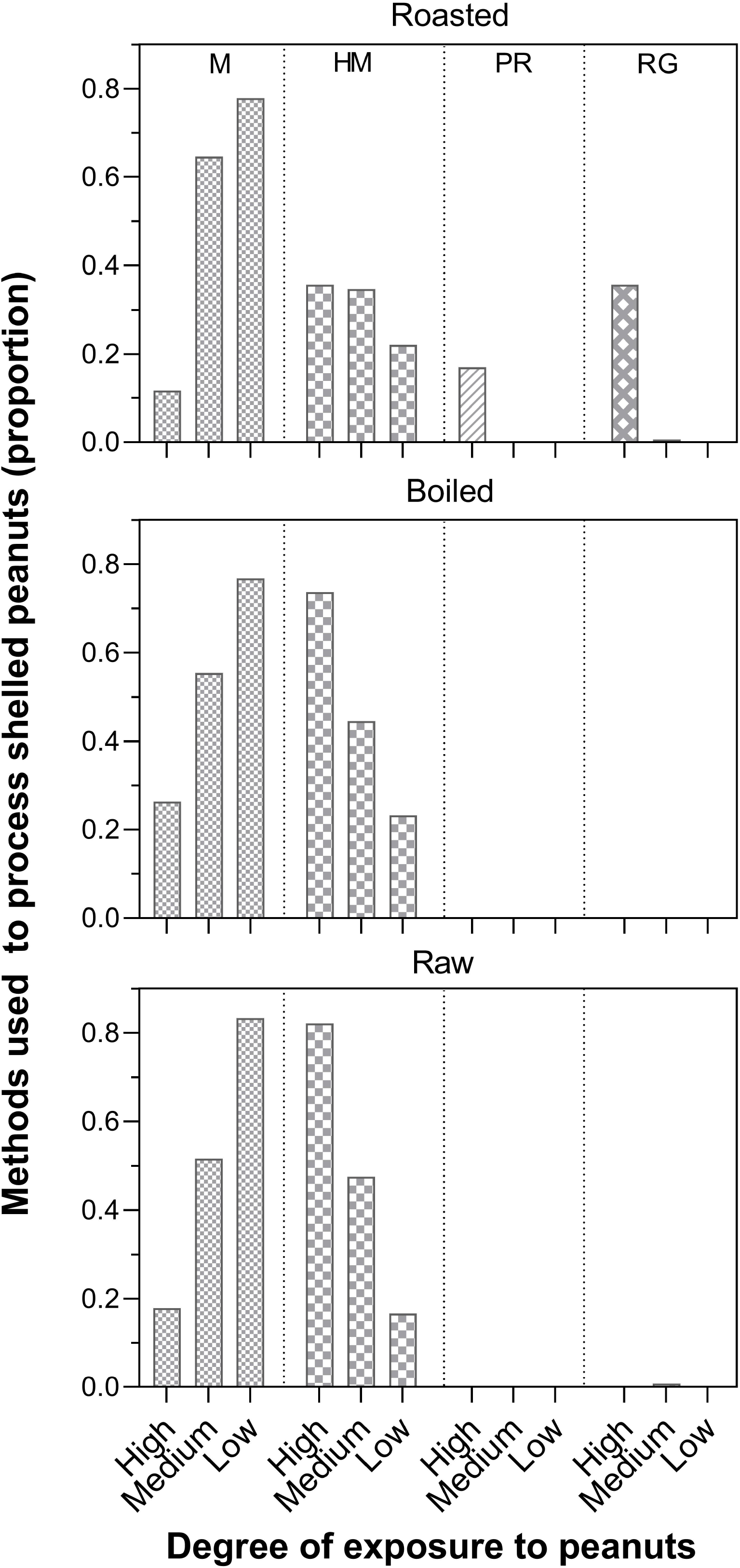
Comparison in the frequency of methods (M-Mouth, HM-Hand and mouth, PR-Palm rub, RG-Rub against ground) used to remove testa of peanut across exposures (High, Moderate and Low) and peanut states (Raw, Boiled and Roasted). The methods adopted are represented in 4 different panels delineated by dotted lines. Bars within a single panel are shaded similarly. Absence of bars represents no data. Alphabets at the top of the bars depict statistical comparison (p<0.05) across exposure types within a method category.

Of the 16 individuals tested in ‘High’ exposure group, we shortlisted 12 individuals with a threshold of ≥6 cases in extraction of roasted nuts. Almost all the individuals flexibly used more than one method for testa removal. The use of HM was the most widespread (ID_HM_=11) followed by RG (ID_RG_=10) and M (ID_M_=8) with PR being concentrated among just 6 individuals. Within the 10 individuals using RG, KA 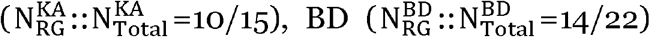 and 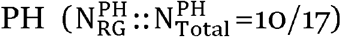 used the extraction method more than once whereas 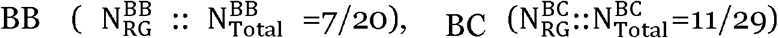, and 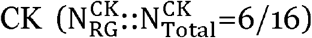 used it at least once (Fig. 3). On the other hand, PR was predominantly used by 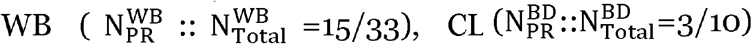, and 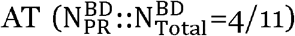 Two older females, CG and CK used allthe recorded methods while 5 individuals (WB, PH, BB, BC, KA and AT) employed three methods to different extents (Fig. 2).

**FIGURE 2.**
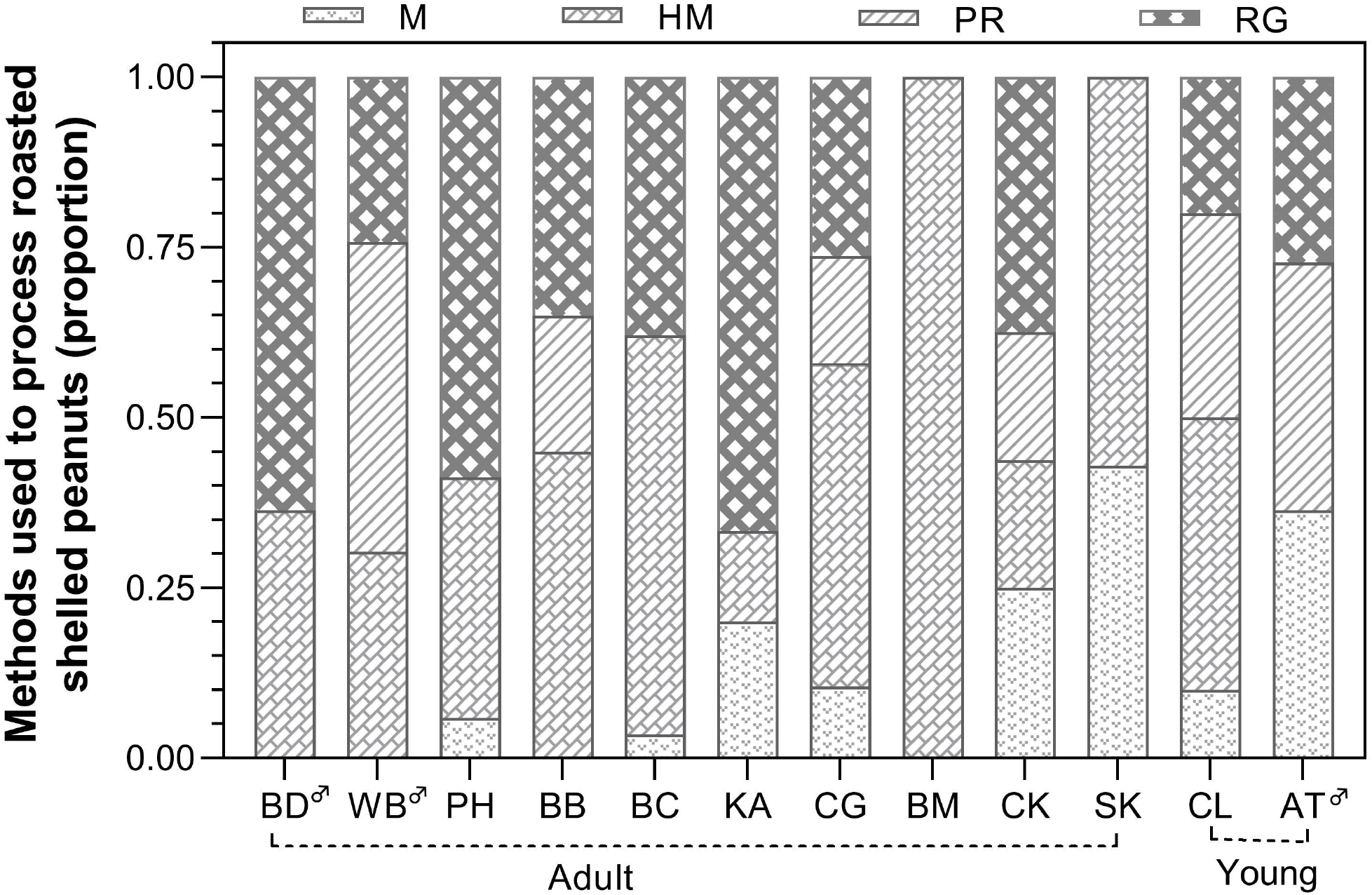
Representation of methods used by select macaques of ‘High’ exposure group to remove testa of shelled-roasted peanuts. Male individuals are identified by the male symbol in the superscript form.

**FIGURE 3.**
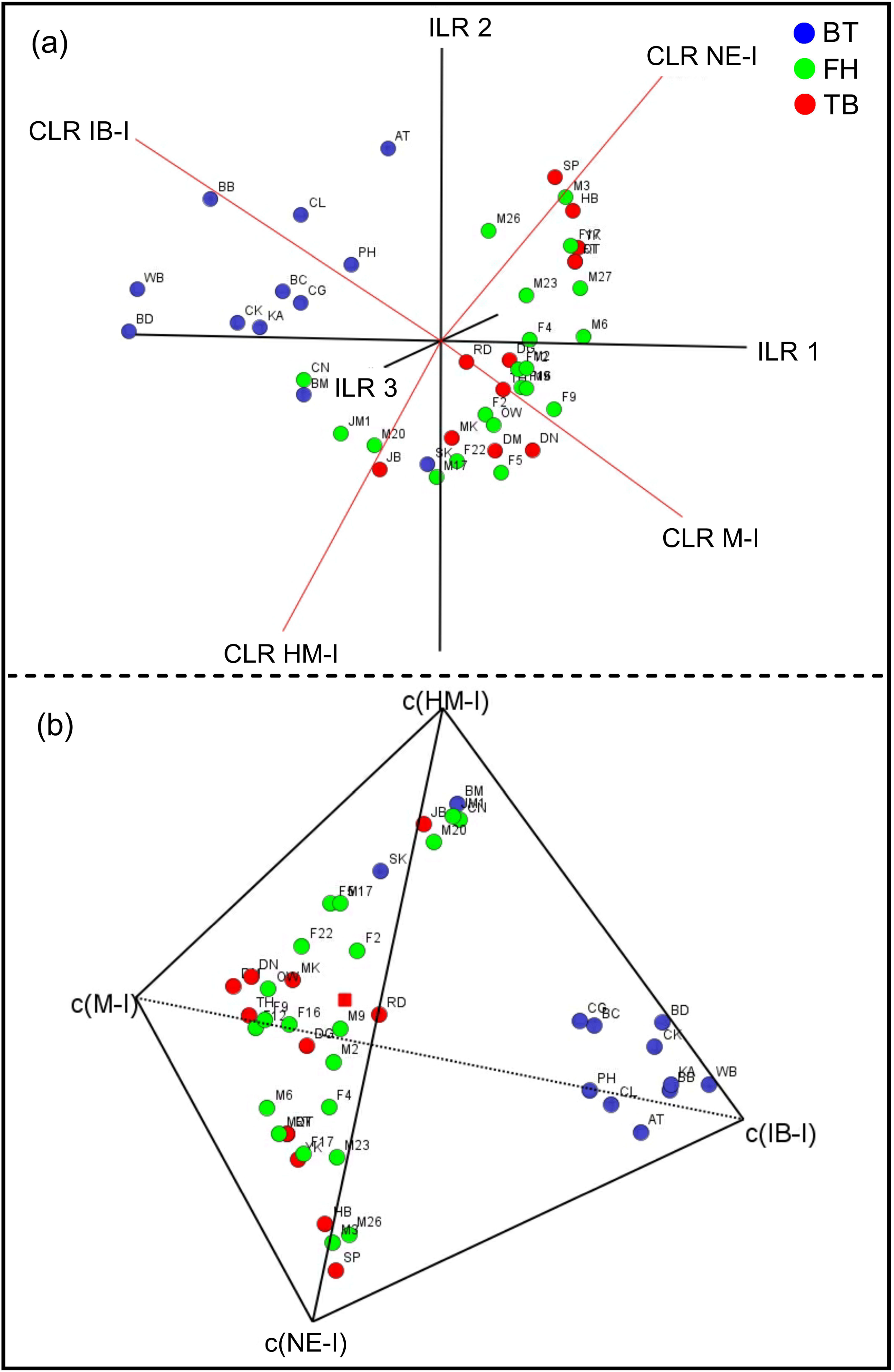
Centered log-ratio (CLR) transformation form Biplot (a) and Quaternary plot (b) of macaques across three groups (BT, FH and TB) based on the relative frequency of non-extraction of (shelled-roasted) peanuts and use of Mouth (M), Hand-Mouth (HM) and Innovative behavior (IB). The center of the sample in (b) is shown as red square. ILR refers to isometric log-ratio transformation; the suffix ‘-I’ signifies imputation by Bayesian multiplicative zero replacement transformation followed by closure; c() denotes centering of compositions.

The compositional data analysis used instances of NE (no extraction) and the methods used in peanut extraction, M, HM and IB, and included macaques ≥ 12 representations across compositional parts (i.e.∑ NE, M, HM, IB). In partial fulfillment of our hypothesis, individuals of BT (N_BT_=13; High exposure) showed a near complete separation from FH (N_FH_=20; Moderate exposure) and TB (N_FH_=11; Low exposure), whereas FH and TB overlapped considerably with each other as represented by their 3D CLR biplot and 3D quaternary plot (Fig. 3). Although unrelated to our hypotheses, visual inspection of three bands of confidence levels (0.90, 0.95 and 0.99) in the four ternary plots (Fig. S4; ^4^C_3_ combinations) showed that the degree of variation within a group varied among the three groups and BT had the biggest area under 99% confidence level across the ternary plots (except in S4(b) without IB as an axis) followed by FH and TB.

### Characterizing methods of testa removal based on duration

Apart from the apparent qualitative distinction among the four methods of testa removal, we found distinct difference in their duration (Dur) of execution. The analysis excluded instances of RG in the ‘Moderate’ exposure category due to a paucity of cases. The mean value of inter-rater consistency in the measure of duration was 0.96 (ICC(2,2); 95% CI: 0.94, 0.98; F=26.62, p<0.0001). Overall, while M (2.98±0.27SE s,N_M_=49) did not differ from HM (4.01±0.33SE s, N_HM_=64) (Dur_M_=Dur_HM_, 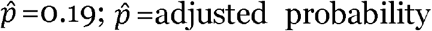) and PR (1.05±0.12SE s, N_PR_=13) did not differ from RG (0.74±0.11SE seconds, N_RG_=22) (Dur_PR_=Dur_RG_,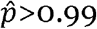) in duration, both PR and RG were quicker than M and HM (K-W=75.39, P<0.0001;Dur_HM_>Dur_PR_, 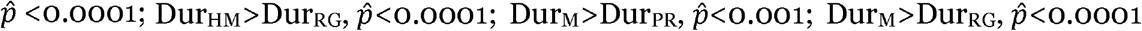). Even when exposure to peanuts was considered, duration of M and HM was comparable across exposures (Dur_M,High_ = Dur_M,Moderate_ = Dur_M,Low_, K-W=5.99, N=3, p=0.05; Dur_HM,High_ = Dur_HM,Moderate_ = Dur_HM,Low_, K-W=3.51, N=3, p=0.17, Fig. 5) and were higher than PR and RG ((Dur_M,All_=Dur_HM,All_)>(Dur_PR_=Dur_RG_); K-W=81.49, N=8, p<0.0001; Fig. 4). We found PR and RG to be 3-4 times and 4-5 times faster than M-HM, respectively. We obtained identical results with multilevel generalized mixed model that controlled for random effects of repeated samples and considered age and exposure as fixed effects to model duration of peanut extraction methods (Table S1; see Fig. S5).

**FIGURE 4.**
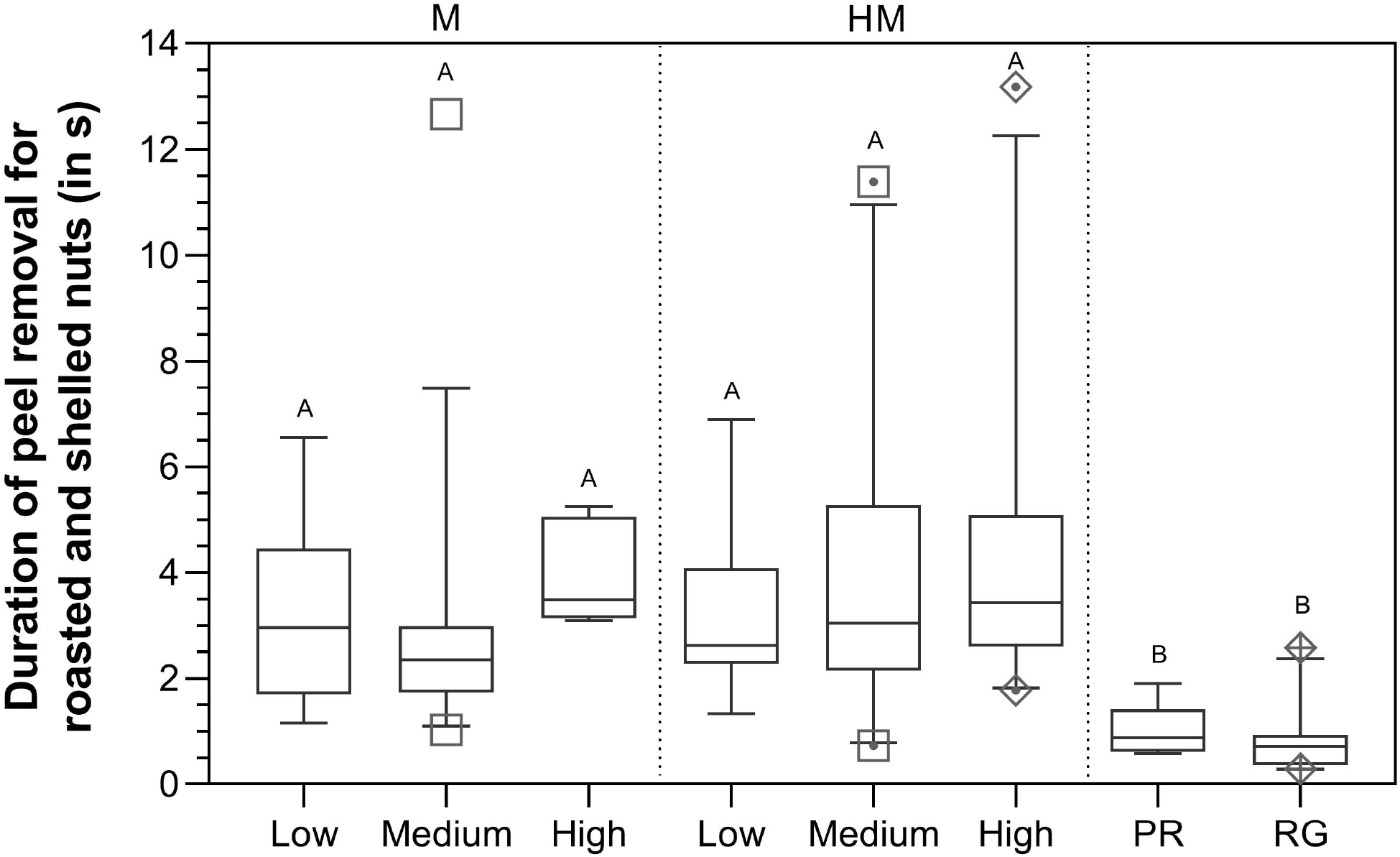
Comparison of the duration of time taken to execute the four methods of testa removal (M-Mouth, HM-Hand and mouth, PR-Palm rub, RG-Rub against ground) across the exposures (High, Moderate and Low). The box-and-whiskers plot stretches from 5-95 percentile and demarcates outliers. The methods of peanut extraction, M, HM and PR-RG are delineated by dotted lines. Palm rub and RG are represented in a single panel since the 2 methods were almost exclusively recorded in ‘High’ exposure category. Symbols used in the figure are unique to sub-categories though blank symbols are used in M and dotted symbols are used inHM. Alphabets at the top of the bars depict statistical comparison (α=95%) across both, exposure types and methods.

### Generalized method of testa removal based on food categorization (Categorization experiment)

We tested nine individuals from the ‘High’ exposure category in the categorization experiment using peas and chickpeas. Of the 69 cases, RG (N_RG_=31) was the most prevalent method of testa removal followed by HM (N_HM_=28), PR (N_PR_=6) and M (N_M_=4). All individuals maintained extraction methods identical to peanuts, albeit WB, who minimized the use of PR (χ^2^=5.79, df=1, p=0.02) in the categorization experiment (Fig. S6). All methods used by a macaque during introductory experiment were not recorded in the categorization experiment due to a low sample size. All incidences of extracting peas and chickpeas using mouth (N_M_=4) in the categorization experiment were solely shown by an adult female, CK.

## DISCUSSION

Using primates as a model, we simulated exposure to processed food in three physically-processed forms and three chemically-processed states to study how extraction decisions differ by exposure to food. The experimental design also offered insight into how even minor treatments of native/natural food could alter its perception and treatment by animals and hence, how essential the ability to categorize food and generalize extractive foraging is for urban synanthropic animals. Ingestion of peanuts in all presentations among all demographics and exposures regardless of their form and state indicated basal familiarity with the physical properties of the nut which were perhaps consequences of previous exposures or conceptual clustering based on similar food items (see Rakison & Oakes, 2009). Nonetheless, exposure-specific latencies to accept certain form(s) and state(s) of peanuts established the degree of familiarity among groups. The process of extracting nuts from native form was remarkably identical regardless of peanut form/state and exposure, suggesting a conserved motor sequence of peanut extraction associated with basal peanut familiarity (cf. with vervet monkeys in Canteloup et al., 2021). In contrast, peanut extraction methods tended towards complexity and optimality with increment in exposure but only for the form and the state of peanut that was the most familiar. Intriguingly, application of the complex methods generalized to food items of similar characteristics via object categorization.

### Familiarity with states and forms of peanuts

Response latency is a veritable measure of memory in cognitive tests (Jahn-Eimermacher et al., 2011) and is routinely used to measure object familiarity (e.g. Anderson et al., 2008; Forss et al., 2015). All groups of bonnet macaques were familiar with the food item albeit to different degree and contingent on its processing status. Desirously, roasting and boiling of raw peanuts only altered the coloration of testa to red and to white, respectively. As a consequence, we did not anticipate a stark attenuation in familiarity but only subtle effect in degree of familiarity that would be measurable through latency. Results of latency of accepting peanuts suggest that familiarity towards peanuts is acquired prior to 2.5 years of age in bonnet macaques. Expectedly, as a result of limited exposure, the ‘Low’ exposure group was relatively the least familiar to peanuts and had a partial familiarity to the boiled state (relative to other states). As opposed to the ‘Low’ exposure group, adequate regular exposure facilitated comparable familiarity of all states of peanuts in the remaining groups. Ironically, despite their familiarity with native and shelled forms, the groups differed strikingly in their familiarity to peeled form. Variability in familiarity of peeled form indicates inherent differences in the final conditions (peel intact/removed) in which peanuts are consumed by the groups.

### Occurrence of testa removal

Age of macaques did not affect occurrence of peanut extraction from its shelled form since all the exposure categories had a basal-level of exposure/familiarity to peanuts and only older immature individuals (>2.5 years of age) could be included in the experiment. The raw state of shelled peanuts attenuated testa removal, while boiled and roasted states prompted depeeling despite the testa in these states being perfectly edible and nutritious (see Yu et al., 2005). Probable explanations for state-specific testa removal could be attributed to the differential mouth feel of testa or alternatively, to the firmness of attachment of the testa to the nut post-processing.

### Methods of testa removal

It is intriguing how a single method of shell removal was used predominantly across exposure while three additional methods varying in motor and cognitive complexities emerged for depeeling peanuts. The ‘High’ exposure group used the most complex extraction methods (PR and RG) equivalently across sex and age albeit in a non-exclusive manner. Conversely, group with the least exposure displayed the highest tendency to use the least complex method (i.e. mouth) of depeeling. The propensity to use mouth and hand-mouth on the basis of the state of the peanut was pronounced in the ‘High’ and ‘Moderate’ exposure categories and equivalent across all states in the ‘Low’ category denoting either an absence of variation or limited familiarity with the states in the latter group. Differential decision of peanut extraction and relative use of extraction methods distinguished BT from the remaining groups. Despite distinct difference in familiarity and exposure between FH and TB, the groups resembled each other closely. Unexpectedly, the degree of inter-individual variation in the use/non-use of extraction methods was also the highest in BT whereas FH and TB were made up of relatively homogeneous individuals.

Innovative behavior often emerges from existing behavioral repertoire of a species for solving novel challenges in which a behavior can either undergo subtle modification or get expressed in a novel situation (Kummer & Goodall, 1985; Ramsey et al., 2007; Reader & Laland, 2003). Palm-rubbing (PR, category-3 manipulation complexity, Guttman, 1944) for the purpose of peanut extraction is an example of the latter kind, wherein ‘food cleaning’, which involves the use of palms and hands to remove dirt and is an ubiquitous behavior among nonhuman primates (bonnet macaque, Roonwol & Mohnot, 1977; Sinha, 2001; bonobo, Allritz et al., 2013; Case et al., 2020; chimpanzee, Allritz et al., 2013; Case et al., 2020; Japanese macaque, Fiore et al., 2020; Suzuki, 1965; gorilla, Allritz et al., 2013; Case et al., 2020; Neadle et al., 2017; Nicobar long-tailed macaque, Pal et al., 2018; Visalberghi & Fragaszy, 1990; Wheatley, 1988; orangutan, Allritz et al., 2013; capuchin, Urbani, 2001; Visalberghi & Fragaszy,1990; vervet monkey, Waal et al., 2012) is utilized to deskin shelled peanut. On the other hand, RG (Category-1 manipulation complexity, Guttman, 1944) aligns with the former characterization of innovation in which ‘food scrubbing’ is modified into rolling against horizontal substrata. Although the indispensability of substratum for undertaking RG is apparent by definition, while establishing statistical validity of the association we found that the presence of substratum is a necessary but an insufficient condition for the expression of RG. Conclusively, determinants of the innovative behavior used in the study are inadequate to explain these extractive foraging behaviors in bonnet macaques. Regardless, in congruence to the theory of ‘Zone of Latent Solutions’ as described by Tennie et al. (2009), we contend that rate of exposure and familiarity to peanuts has driven the expression of latent solutions (extraction behavior) in the form of innovated/re-innovated foraging extraction in macaques.

### Benefits and demands of the innovative behavior

The emergence of the innovative methods of peanut extraction can be explained by their advantage over existing methods. Although mouth use and hand-mouth use efficiently depeeled nuts, both the innovative methods were much quicker. Thus, the use of PR and RG has the benefit of accelerating feeding than is possible with the remaining methods but with an additional requirement of an artificial surface for RG. In fact, naturalistic observations of PR and RG were often associated with a large number of scattered shelled-roasted peanuts. However, PR and RG require advanced perceptual-motor and sensory-motor skills that comprise of the following steps: 1) Orienting nut (PR: on palm; RG: on horizontal smooth substratum) along the saggital plane mediated by visual cues and static tactile cues, 2) positioning processing hand (PR: non-contact hand over the other hand; RG: processing hand over substratum) such that the nut is in contact with the palm(s) controlled by static tactile cues, 3) rubbing nuts (PR: against palms; RG: rolling nuts against substratum) through perception of dynamic pressure and dynamic tactile cues followed by a cognitive prediction of the nut condition and 4) cessation of rubbing/rolling based on either, visual/auditory/tactile cue of the separation of testa or cognitive prediction. As a result, we found young individuals struggle to use PR and RG, which denoted an ontogenetic constraint in the use of these methods. ‘Rub on ground’ was the most widespread method among the two innovative behavior prevalent in over 2/3^rd^ of BT. In comparison, PR was limited to only six individuals possibly due to the demand of high sensorimotor and coordination skills, often requiring multiple attempts for successful extraction.

### Generalization in method of testa removal

The successful extension of innovative extraction to peas and chickpeas by BT is symptomatic of the extraordinary ability of urban synanthropic groups to engage in category learning. From a sensory perspective, categorization and/or conceptual clustering might have considered common physical properties of roasted peas, chickpeas and peanuts, for instance, flaky skin and near spheroidal shapes (peanut, Wanget et al., 2019; pea, Zhao et al., 2020; chickpea, Wood et al., 2011). While, flaky testa of (roasted) pea and (roasted) chickpea stimulated de-peeling, spheroidal shape facilitated rubbing and rolling actions during PR and RG, respectively. Co-opting or general application of the innovative extractive foraging behavior based on food categorization further suggests a profound cognitive understanding of the techniques in the urban macaques.

As urbanization drives biotic homogenization globally (Dar & Reshi, 2014; M. L. McKinney, 2006), we view our study to be a small step towards building a comprehensive research on how synanthropic/synurbic species make dietary and extraction decisions in the vast and dynamic foraging landscape of processed/synthetic food in perpetually-altering packaging. Apart from the severe long-term effects of maladaptation to processed food consumption in animals, i.e. alopecia (Maréchal et al., 2016), obesity (Klimentidis et al., 2011) and cardiovascular diseases (Zaragoza et al., 2011), the criticality of urban foraging decisions is truly realized in context to deducing edibility of non-natural objects wherein inability/failure may result in poisoning and/or death. However, since we reckon that stimulus-centered experimental designs will ultimately offer requisite insights, research in the field will need to address and contend two pivotal issues: 1) Theseus’s paradox (of a slightly different version), i.e. when do/does the primary ingredient/(s) lose its/their identity/(ies) and assume the identity of the final (processed) product (cf. Morell-Hart, 2020) and 2) ethics surrounding experimentation with non-natural food items. We surmise that this theme of research can contribute profound insights into the urban adaptation of animals, formulate strategies to curtail dependence of animals on processed food and devise plans for the conservation of threatened synanthropic species.

## MATERIALS AND METHODS

### Study groups

Of the 3 moderate-sized groups of bonnet macaques chosen, two (Bull Temple, BT and Foot Hills, FH) of them were altitudinally stratified in Chamundeshwari hills, a major site for religious tourism and the third (Tirumakudal Betahalli, TB) subsisted in an arid agricultural landscape in Mysore, Karnataka. All the study groups have been under intermittent monitoring for circa. 30 years and hence each of their demographic history, feeding ecology, ranging behavior and anthropogenic pressures are qualitatively established (see Singh, 2019).

BT derived its nourishment largely from anthropogenic sources like garbage dumps, food carried/discarded by visitors (like flavored milk-based drinks, fruit drinks, nuts, fruits, flowers, etc.), food carts and human settlements, and spent 7-9 hours per day in anthropogenic spaces. A little less than a fourth of the diet of BT was natural, derived from a dry deciduous forest reserve. Based on McKinney’s (2015) anthropogenic influence classification system, the group was coded as F_3_J_3_G_3_B_2,3_. BT had the highest exposure to all states and forms of peanuts except in the raw state and encountered roasted shelled peanuts the most (independently as well as in other processed food like fried cutlet and sweets).

The second group, FH was as much dependent on anthropogenic food resources as BT, though active food acquisition by snatching and stealing were relatively lower and food provisioning by dedicated visitors were much higher, especially from fruit and vegetable vendors. Few adult macaques were abnormally obese and had strikingly large patches of alopecia (10-25% of body surface) in the group (see Zhang, 2011). We coded FH group as F_3_F_2,3_G_2,3_B_2_. The group had seasonal access to native form of peanuts and accosted roasted shelled peanuts throughout the year but at a lower rate than BT. Qualitatively, FH had a moderate exposure to peanuts.

In contrast to BT and FH, TB foraged on agricultural crops and natural resources and included a small subgroup of macaques that derived a minor portion of nourishment from households. Despite the absence of artificial provisioning, TB encountered large swathes of visitors/devotees once in four years for a period of one week during a religious festival. We coded FH group as L_1,3_H_1,2,3_N_3_N_3_. The only probable occasion for TB to encounter peanut is during household foraging and during the local festival and therefore, the group had the least exposure to peanuts.

### Experimental design

The ‘introductory’ experiment used three forms of peanuts, (1) native (pod intact), (2) shelled (pod removed) and (3) skinned (pod and testa, i.e. seed skin removed), each presented in three states, (1) raw, (2) roasted and (3) boiled. The native and roasted forms were directly acquired from vendors while the boiled states, regardless of form was prepared by boiling raw native peanuts in potable water for approximately 20-30 minutes until the peanuts blanched and softened. The de-skinned form of peanut was used as a control to attest familiarity with peanut and to distinguish extraction methods. The ‘categorization’ experiment was carried out on select individuals and used roasted forms of pea and chickpea, both of which were characterized by flaky and loose skin. A minimum of 2 trials were carried out using pea and chickpea, each.

We targeted isolated and passive macaques while conducting an experimental session. If a target macaque was seldom observed in a solitary state, then the entire social group was offered peanut (but slightly distant from each other such that possibility of social learning is minimized) and the target macaque was recorded. The experimental sessions were scheduled either between 0700-1100h or between 1500-1900h. Within each session, we conducted a maximum of three presentations and maintained a minimum inter-trial interval of 2 minutes during consecutive trials. A gap of at least 30 minutes was maintained between sessions if the same individual was tested. Once an isolated macaque was selected, a researcher/volunteer less familiar to the macaque placed 1-2 units (pod/seed) of peanut on the ground along its direction of vision, and recorded the behavior of the individual either, till each nut was ingested or the individual moved out of sight. An intermediate familiarity (of offeror) allowed achieving a balance between attraction caused by the expectation of food and repulsion of unknown human (see Johnson, 2000). We sampled individuals across demography, reproductive state, social rank, gregariousness and human habituation as per the recommendations of STRANGE framework in behavioral sampling (see Farrar & Ostojić, 2021; cf. Webster & Rutz, 2020). A trial was considered invalid if ingestion of peanuts could not be verified and no specific order was followed while presenting peanuts. We conducted a maximum of three trials with each macaque for each form and state of peanut. Since video recording of participating macaques was possible even at lower proximity (<15m), habituation of the groups to our presence was not necessary.

The experiments were carried out in three phases each lasting for 2-3 months; from September to November, 2014 with the BT group, from May to July, 2015 with the FH group and finally, from January to February, 2019 with the TB group. We used three models of camcorders (Sony DCR-SX21, Sony DCR-DVD 650 and Sony HDR-CX405) to record the experiments, one in each phase of field experimentation.

### Behavioral coding and data analysis

Video recordings of the experiment were coded manually by two researchers (TD and SD). To establish the degree of familiarity with peanut, we measured latency of acceptance (in second), either as the time period between physical contact with the nut and stabilization of extraction (i.e. when visual/olfactory/gustatory exploration ceased) when offered in native/shelled forms or feeding when offered in peeled form in a random subset of experiments. Recordings with obscured approach/movements and instances of food-cleaning prior to extraction were not coded. Latency of acceptance was measured using SPER Scientific 810029 Stopwatch. Each researcher followed the fate of individual nut regardless of form. When presented in native form, we noted occurrence of pod removal, method of pod removal, occurrence of testa removal and method of testa removal. Similarly, when presented in shelled form, occurrence of testa removal and method of testa removal were recorded during introductory and confirmatory experimental phases. In the control scenario where de-skinned nuts were presented, only the occurrence of ingestion was coded. Lastly, durations of extraction methods were determined from initiation to completion of peanut extraction (i.e. removal of testa) in a random subset of experiments with shelled peanuts.

Statistical analyses were performed using RStudio v.1.4.1717 (RStudio Team, 2021). We used multilevel generalized linear mixed modeling to test the main effects of age, exposure, form and state of peanuts and interaction effects of exposure and form/state of peanuts on latency of acceptance. Multilevel generalized mixed modeling was used to determine the influence of state and exposure on removal of peanut skin and methods used to extract peanuts using either, glmer() function in lem4 package (Bates et al., 2015) or glmmTMB() function in glmmTMB package (Brooks et al., 2017). Model diagnostics of the multilevel models was performed using DHARMa package (Hartig, 2021) following Dunn & Smyth (1996) and Gelman & Hill (2007). For testing within-group similarity and inter-group differences in relative use of extraction methods, we used Compositional Data analysis (CoDa; see Greenacre, 2016) since proportion of response to peanut extraction (including non-occurrence of extraction; compositional parts) summed to 1 and hence, was not independent. We used CoDaPack v.3.02.21 (Comas-Cufí & Thió-Henestrosa, 2011) for data analytics and visualized the compositions as CLR (Centered log-ratio transformation) form biplot and ternary plot following Greenacre (2016). Comparison of duration of peanut extraction by each extraction method across exposure categories was done using Kruskal-Wallis test followed by Dunn’s correction for multiple post hoc comparisons.

## ACKNOWLEDGEMENTS

The authors are much obliged to the Executive Officer, Sri Chamundeshwari Temple, police officers at Chamundi Hill Police station, and residents of Pourakarmika colony, Chamundeshwari Bettada Paada and Tirumakudall Betahalli for their magnanimity and in indirectly ensuring unabated field experimentation. Field experiments were carried out with assistance from Mr. Mohan Kamraj, Mr. Manikanta M., Mr. Jeril Joy, Mr. Mohan A., Mr. Shyam Raj, Mrs. Deepa S.M., Mr. Kiran Kumar, Mr. Bharath Kumar, Mrs. Preethi Bharath, Mr. Vinay L., Mr. Bharath Nayak, Mrs. Jayanti Kashyap, Mr. Darshan Sosale, Dr. Charles Sosale, Mrs. Mala Charles, Ms. Divya Sosale, Ms. Kruthi Reddy, Mrs. Parimala Reddy, Ms. Aishwarya Anand and Ms. Apoorva A. S. and we appreciate each of their effort. We deeply appreciate statistical assistances provided by Dr. Swetashree Kolay and Mr. Nisarg Desai. SD expresses his gratitude to University Grants Commission, Government of India for his doctoral fellowship; AKV and PG are grateful to the Summer Research Fellowship Program of the 3 science academies of India; MS is thankful to Science and Engineering Research Board, Government of India sponsored ‘Distinguished Fellowship’ which also supports TD.

## Ethical statement

The study was approved by the Institutional Animal Ethics committee of the University of Mysore and abided by the Code of Best Practices for Field Primatology of the International Primatological Society and the American Society of Primatologists formulated in 2014.

## Data availability

The entire dataset utilized to report this article will be shortly made available in the Dryad repository. The dataset will be linked to this article before publication.

## Supporting Information

Supporting information can be found in the online version of the article under the Supporting Information section.

